# Chasing non-existent “microRNAs” in cancer

**DOI:** 10.1101/2024.12.05.626946

**Authors:** Ayla Orang, Nicholas I Warnock, Melodie Migault, B Kate Dredge, Andrew G Bert, Julie M Bracken, Philip A Gregory, Katherine A Pillman, Gregory J Goodall, Cameron P Bracken

## Abstract

MicroRNAs (miRNAs) are important regulators of gene expression whose dysregulation is widely linked to tumourigenesis, tumour progression and Epithelial-Mesenchymal Transition (EMT), a developmental process that promotes metastasis when inappropriately activated. However, controversy has emerged regarding how many functional miRNAs are encoded in the genome, and to what extent non-regulatory products of RNA degradation have been mis-identified as miRNAs. Central to miRNA function is their capacity to associate with an Argonaute (AGO) protein and form an RNA-Induced Silencing Complex (RISC), which mediates target mRNA suppression. We report that numerous “miRNAs” previously reported in EMT and cancer contexts, are not incorporated into RISC and are not capable of endogenously silencing target genes, despite the fact that hundreds of publications in the cancer field describe their roles. Apparent function can be driven through the expression of artificial miRNA mimics which is not necessarily reflective of any endogenous gene regulatory function. We present biochemical and bioinformatic criteria that can be used to distinguish functional miRNAs from mistakenly annotated RNA fragments.

## Introduction

MicroRNAs (miRNAs) function as the target specificity component of the RNA-induced silencing complex (RISC) [1]. With few exceptions, they are generated by sequential processing of primary transcripts, involving recognition and cleavage of a precursor RNA in the nucleus by the endonuclease Drosha to generate a hairpin pre-miR, followed by cleavage in the cytoplasm by Dicer, producing a linear RNA of close to 22 nt that is loaded into the Argonaute (AGO) proteins. In humans there are 4 AGO proteins that share high sequence similarity, with AGO2 being the most abundant in most tissues [2]. MicroRNAs act as the target recognition sequence to bind AGO to the 3’UTR of mRNAs, primarily through base pairing of the “seed region”, comprising nucleotides 2-8 at the 5’ end of the miRNA [3].

MicroRNAs have been found to affect many aspects of cell differentiation and function, and also to act as oncogenes or tumour suppressors in many types of cancer [4, 5]. They are prominent in the control of epithelial to mesenchymal transition (EMT) [6, 7], which has implications for cancer metastasis and chemoresistance [8]. However, questions have emerged regarding the validity of the identification of many small RNAs that have been given the designation of “microRNA” [9–14]. For example, while the online sequence repository miRBase (http://www.miRBase.org), lists over 2700 human miRNAs [15], the more stringent repository MirGeneDB lists less than 600 high confidence human miRNAs [16].

In this study, we identify dozens of miRbase-listed small RNAs that do not bind AGO and hence, do not function as miRNAs. This is despite hundreds of publications in which their roles in EMT and/or cancer have been reported. We compare the features of these non-miRNAs to those of genuine miRNAs, describe how the mis-interpretation of exogenous expression experiments are likely responsible for these contradictory findings and provide a series of experimental and bioinformatic criteria that can be used to help distinguish functional miRNAs from adventitious RNA fragments that have been mistaken for miRNAs.

## Materials and Methods

### Cell culture and treatments

The Human Mammary Epithelial Cells (HMLE) cell line was obtained from the Weinberg lab [17] and grown in HuMEC Ready Media (HuMEC Basal Serum-Free Medium (Gibco, #12753018) supplemented with HuMEC Supplement Kit (Gibco, #12755013)). A HMLE subline with a mesenchymal phenotype (mesHMLE) was achieved by culturing in DMEM:F12 media (Gibco, #11320033) supplemented with 10 mg/ml insulin (Actrapid HM 100IU/ml Penfill), 20 ng/ml EGF (R&D systems, #RDS236EG01M), 0.5 mg/ml hydrocortisone (Sigma Aldrich, # H0888), and 5% fetal Bovine serum (Hyclone, #SH300084) and treatment with 2.5 ng/ml of recombinant Human TGF-β 1 Protein (R&D systems, #RDS240B002) for maximum of 18 days. MesHMLEs were then maintained in this supplemented media without TGFβ1. Cells were grown at 37°C with 5% CO2.

### Real-time PCR

After RNA quantification using an ND-1000 NanoDrop spectrometer, 1 µg of RNA was reverse transcribed into cDNA using the QuantiTect Reverse Transcription Kit (Qiagen, # 205311). Real-time polymerase chain reactions were performed on a Corbett Rotor-Gene 6000 Detection System (Qiagen) using a QuantiTect SYBR Green PCR Kit (Qiagen, # 204141). For quantification of small RNAs, multiplex TaqMan MicroRNA Reverse Transcription Kit (Applied Biosystems, #4366596) and 2x Universal PCR Master Mix (Applied Biosystems, #4324018) were used following manufacturer’s instructions with assay IDs as follow: U6snRNA #001973, hsa-miR-16-5p #000391 and miR-222-3p #002276.

### Subcellular fractionation and RNA extraction

Subcellular fractionation was carried out as described previously [18]. Briefly, confluent cells in a T75 flask were trypsinized and collected in a 15 mL conical tube, washed once with ice cold 1XPBS (Gibco, #70011044) and resuspended in 400 µL of Hypotonic Lysis Buffer (HLB: 10 mM Tris pH 7.5 (Invitrogen , #15504020), 10 mM NaCl (chem Supply, # SA046), 3 mM MgCl_2_ (Merck, #7791186), 0.3% (vol/vol) NP-40 alternative (Calbiochem, #492016) and 10% (vol/vol) glycerol (Univar Solutions, #242), supplemented with 100 U of SUPERase-In (ThermoFisher, # AM2694) and left on ice for 10 min. Nuclei were separated by centrifugation at 1000 g for 3 min and the supernatant was collected as the cytoplasmic fraction. The nuclear pellet was then washed four times in 1 mL HLB buffer and each time pelleted by centrifugation at 200g for 2 min. 1 mL RNA Precipitation Solution (0.5 ml of 3 M sodium acetate pH 5.5 (AnalaR, #10236)) in 9.5 ml of ethanol (Univar Solutions, #AJA214) added to the cytoplasmic fractions and incubated at -20 °C for 2 hr before centrifugation at 18000 g for 15 min. The cytoplasmic and nuclear pellets were then resuspended in Trizol Reagent (Ambion, #15596018) and RNA extraction was carried out according to the manufacturer’s protocol.

### AGO:RNA co-immunoprecipitation

HMLE and mesHMLE cells were grown in 10 cm dishes (∼1 dish per IP), harvested by scraping in ice-cold PBS and centrifugation at 500 g for 5 mins and pellets stored at –80°C until use. Cell pellets were lysed on ice for 10 min using 1mL lysis buffer (1X PXL) containing 1X PBS, 0.1% SDS (Sigma Aldrich, #75746), 0.5% deoxycholate (Sigma Aldrich, #D6750), 0.5% NP-40 (IGEPAL, #CA-630) supplemented with protease inhibitors (Roche, #4693159001) and RNAseOUT (Invitrogen, #10777019) and triturated by passing through a 21G needle. Lysates were treated with 10 µl TurboDNase (Ambion, #AM2238) at 37°C for 10 min, 400 rpm in a Thermomixer then placed on ice >1min. Ethylenediaminetetraacetic acid (EDTA-Univar Solutions, #A180) was then added to a final concentration of 30 mM and lysates centrifuged at 35000 rpm for 20 mins in a Beckman TLA55 ultracentrifuge at 4°C. 350 µg clarified lysate at 0.4 mg/ml was then subjected to immunoprecipitation for 75 min with a pan-anti-Ago antibody (4F9,[19]) pre-bound to Pierce protein-L magnetic beads (Thermo Scientific, #88849); 21 µg Ab, 20 µl beads per IP. Beads were washed twice each with ice-cold 1X PXL, 5X PXL (5X PBS, 0.1% SDS, 0.5% deoxycholate, 0.5% NP-40), and 1X PNK (50 mM Tris pH 7.5, 10 mM MgCl_2_, 0.5% NP-40) prior to digestion with 2 mg/ml proteinase K (Invitrogen, #25530049), 50°C, 1hr. RNA was extracted using Trizol according to the manufacturer’s instructions followed by treatment with TurboDNase (Ambion, #AM2238), phenol (Ambion, #AM9712) extraction and ethanol precipitation. Small RNA libraries were prepared using NEXTflex Small RNA-Seq Kit v3 (Perkin Elmer, #NOVA-5132-05) with 17-18 PCR amplification cycles, size purified by PAGE , extracted by the “crush and soak” method as previously described [20] and sequenced on an Illumina NextSeq 500 (1 x 75 bp).

### Small RNAseq

For fractionated samples RNA was extracted according to Trizol manufacturer instructions. The quantity and quality of the RNA samples were checked using nanodrop Qubit HS RNA (Invitrogen, # Q32852) and Bioanalyzer small RNA assay (Agilent, #5067-1548). Libraries were generated using 1 µg of the total RNA via QIAseq miRNA kit (Qiagen, # 331502) with 14 cycles of amplification. Amplified barcoded libraries were then size selected using auto gel-purification Pippin prep 3% agarose (SAGE science) which targets ranges from 100-250 bp. Libraries (size between 180-190 bp) were then confirmed by Qubit HS DNA and Bioanalyzer HS DNA assay for size and concentration. The libraries were then pooled together in equimolar amounts and sequenced using an Illumina Nextseq 500 using a 1x75 cycle high output kit.

### Reporter construct design and cloning

PsiCHECK2 dual luciferase reporters (Promega, #C8021) were constructed to contain a single copy of a perfectly complementary target site of the small RNA sequences derived from miRBase. Oligonucleotides (IDT technologies) were designed with XhoI (NewEngland BioLabs, # R0146S) and NotI-HF (NewEngland BioLabs, # R3189S) overhangs (Table S1), annealed and ligated into psiCHECK2 using XhoI and NotI restriction sites.

### Luciferase reporter assays

HMLE and mesHMLE cells were seeded in 96-well plates (10,000 cells/well) and incubated overnight at 37°C/5% CO_2_. Before transfection, media was changed, and cells were co-transfected with 5 ng of miRNA psiCheck2 reporter constructs and 5 nM of miRNA mimics or 20 nM of miRNA inhibitors (GenePharma) with Lipofectamine™ 2000 Transfection Reagent (Invitrogen, #11668019), according to the manufacturer’s protocol. Media was changed 6 h post-transfection. After 48 h of transfection, luciferase activity assay was performed using Dual-Luciferase® Reporter Assay System (Promega, #E1980) on a GloMax®-Multi Detection System (Promega), according to the manufacturer’s protocol. Relative luciferase activity was calculated as the ratio of Renilla to Firefly luciferase activity. The Firefly luciferase gene expressed from the same vector was used as an internal reference and cells transfected with empty psiCHECK2 vector were used as control. All reporter assays were performed as at least 4 technical replicates (individual wells) per experiment, with the entire experiment repeated independently. Box plots represent median and interquartile range and whiskers, the minimum and maximum values. miRNA mimics and inhibitors were provided by GenePharma as double-stranded and single stranded RNAs, respectively (Table S2).

### Bioinformatic analyses

FASTQ files from cellular fractionation experiments were analysed at various stages for quality and content using FastQC v0.11.9 [21]. Raw reads were adapter trimmed and filtered using cutadapt v2.8 [22] using an adapter sequence of AGATCGGAAGAGCACACGTCTGAACTCCAGTCAC, minimum length of 28, error rate of 0.2, and overlap of 5. Reads derived from PCR duplication were collapsed using Unique Molecular Identifiers (UMIs) using UMI-tools (v1.0.1; [23]) by first using the ‘extract’ method to cut the UMIs from the 3’ end of the reads. Next, a 3’ spacer sequence was removed using cutadapt (adapter: AACTGTAGGCACCATCAAT, minimum length: 18, error rate: 0.2, overlap: 5). The resulting reads were mapped against the human reference genome (build hg19) using the BWA v0.7.17 alignment algorithm [24] with default parameters, resulting in an average alignment rate of ∼84%. Subsequently, PCR duplicate reads were collapsed using the UMI-tools ‘dedup’ method (using -- method=percentile for practicable memory usage). Alignments were visualized and interrogated using the Integrative Genomics Viewer v2.8.0 [25]. RNA classes were analysed using GRCh37 annotation of gene biotypes from download from Ensembl [26].

Annotation obtained from miRBase (www.miRBase.org; [27] was used to obtain miR-level read counts using the python scripts described in [28], with miRNA-derived reads comprising an average of ∼22% and ∼15% of the nuclear and cytoplasmic fractions, respectively. To compare miRNA abundances between the nuclear and cytoplasmic fractions, miRNA-level library sizes from cellular fractionation experiments were TMM-normalised [29] using the calcNormFactors function from the edgeR package [30] and counts expressed as log counts-per-million (log cpm; log base 2, prior count of 0.1). The mean of biological replicates was used to calculate log2 ratios by subtracting the cytoplasmic value from the nuclear value. The same approach was used to produce a log2(AGO-IP:smallRNA) ratio for each annotated miRNA in the AGO:RNA co-immunoprecipitation small RNA sequence data.

To study the distribution of miRNA read lengths, isomiR-level library sizes were TMM-normalised and counts expressed as cpm. The mean cpm of each isomiR was calculated across biological replicates, and values from isomiRs of common length belonging to each miR were summed for visualisation.

## Results

Annotated “microRNAs” that are implicated in cancer but not associated with Argonaute

The scientific literature is replete with reports of miRNAs that affect EMT, suggesting that many different miRNAs are involved in regulating EMT or in mediating its downstream consequences [6]. However, in the past, some miRNA-sized RNAs may have been erroneously designated as miRNAs, leading us to ask whether all the reports of EMT-regulating miRNAs are reliable [31]. To address this, we examined the changes in miRNA content of RISC during EMT, by sequencing the small RNAs that immunoprecipitate with AGO in HMLE immortalised human mammary epithelial cells and in their TGF-β-induced mesenchymal derivative, mesHMLE cells (Fig. S1). Considering only “miRNAs” that are expressed at more than 5 counts per million in these cells, we found that 7% of the annotated “miRNAs” in HMLE and mesHMLE cells are not actually bound to AGO, making it unlikely they can function as miRNAs (Fig. 1A and Fig. S1, Supplementary Table 3). This includes 26 miRNAs that have been reportedly involved in EMT and/or a cancer context, with more than 250 papers that name the “miRNA” in the paper title (Fig. 1B).

**Figure 1.**
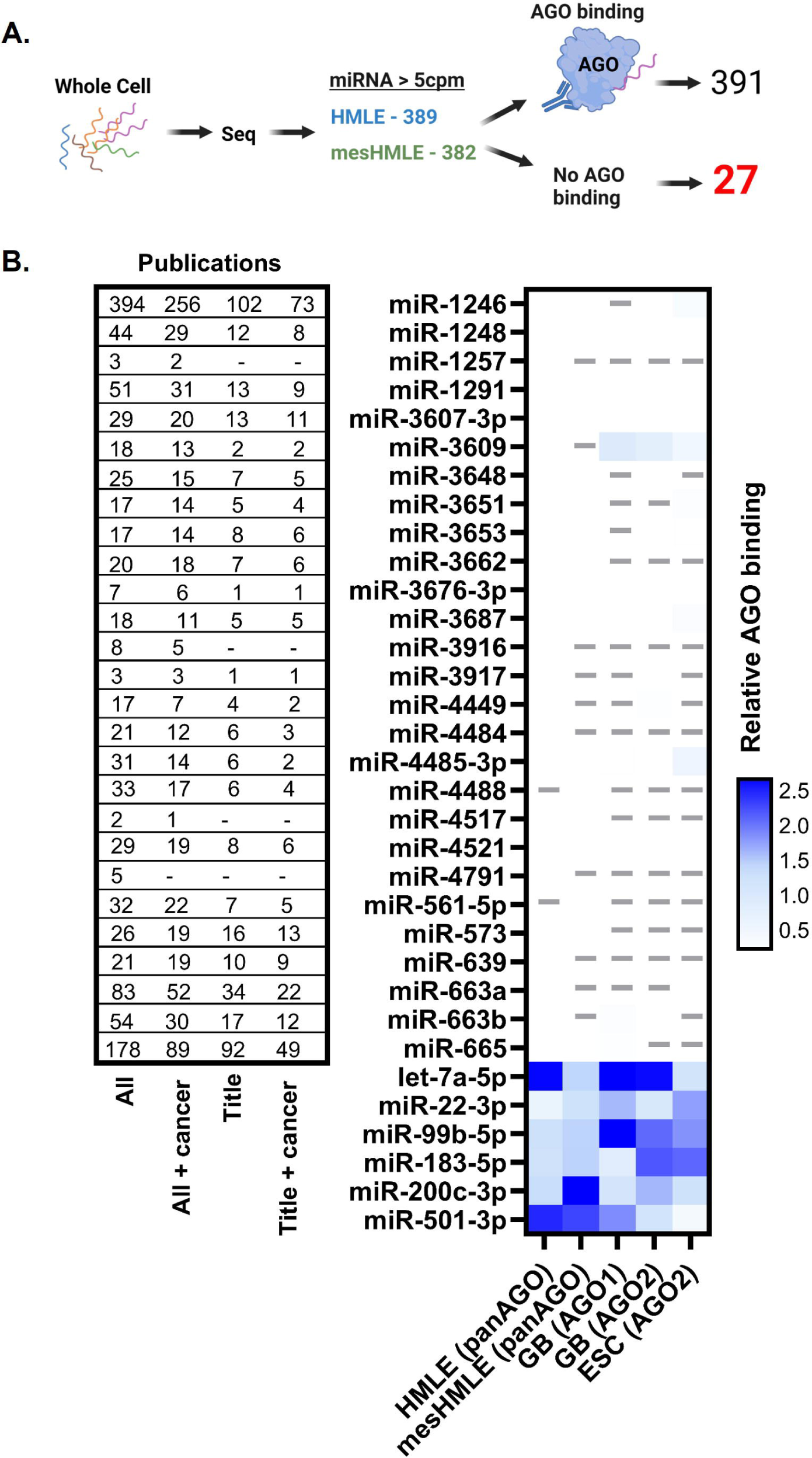
A subset of annotated microRNAs do not bind Argonaute. (A) 418 microRNAs are expressed at >5 counts per million in HMLE and/or mesHMLE cells. 27 of these display no AGO-binding capacity. (B) AGO binding efficiency (calculated by ratio of read counts mapping to each miRNA in the sequencing of the AGO co-precipitate and whole cells), is represented by heat map (darker blue equates to more efficient AGO binding, the final six miRNAs are included as positive controls, representing relative AGO binding of well established, unequivocal miRNAs). Data are drawn from our HMLE / mesHMLE cells performing the Ago precipitation with a pan-AGO antibody, and from publicly available data [32] in which AGO1 or AGO2-specific antibodies were used to immunoprecipitate AGO in glioblastoma and embryonic stem cells. Dashes indicate insufficient miRNA expression to determine AGO binding capacity. The table indicates the number of publications (Pubmed) whereby the miRNA in question features in the title or abstract (“all”) or specifically the title, with or without “cancer” as a co-specified search term.

To assess whether these “miRNAs” are similarly not bound to AGO in other biological contexts, we examined publicly available matched whole-cell RNA sequencing and AGO-coprecipitation data sets from glioblastoma and embryonic stem cells [32]. None of these miRNAs were associated with AGO in any of those datasets (Fig. 1B), making it unlikely they really are functional miRNAs. Hereafter we refer to these miRNA-sized RNAs that have been reported as miRNAs but do not associate with AGO as “non-miRs”.

### Small RNAs that do not bind AGO do not repress reporter genes

To confirm that the non-miRs we had identified do not function as miRNAs, we tested the capacity of several to inhibit target mRNAs despite their lack of association with AGO. We made luciferase reporter genes with perfect matches to 5 of the non-miRs (which collectively feature in over 200 publications). As positive controls, reporters were made for 7 authentic miRNAs that we had confirmed to be associated with AGO. We also made reporters for two arbitrary RNA sequences that do not exist in the genome. To check the reporters can respond to each RNA if they are ectopically presented as miRNA mimics, we co-transfected each luciferase reporter with its corresponding miRNA mimic and saw that there was strong repression of luciferase in each case, including the controls (NT-miR and NT-miR-5U) that are not present in the genome (Fig. 2A). Then, to test whether the endogenous non-miRNAs actually inhibit the reporter when not overexpressed as miRNA mimics, we asked whether antisense inhibitors relieve repression of the reporter, thereby leading to increased luciferase activity. While antisense inhibition of each of the positive control miRNAs increased reporter expression as expected, there was no increase in reporter expression by antisense inhibition of any of the non-miRNAs, either in HMLE cells or in mesHMLE cells (Fig. 2B-C, Supplementary Figure 2). The capacity of an antagomir to de-repress an endogenous luciferase reporter should be commensurate with the level of endogenous miRNA expression. In almost all cases in HMLE cells, and with miR-3607 in mesHMLEs, non-miR antagomiRs failed to de-repress the reporter, despite the non-miR being expressed at levels equal to or greater than genuine miRNAs whose inhibition did increase reporter activity (Fig. 2D-E). This confirms that small RNAs that are not resident in the RISC complex do not function as miRNAs. It also demonstrates that ectopic expression of miRNA mimics (as in Fig. 2A) does not provide evidence that an endogenous small RNA is a miRNA. Instead, it demonstrates that gene-suppressive functions can be artificially engendered by transfection with artificial miR mimics.

**Figure 2.**
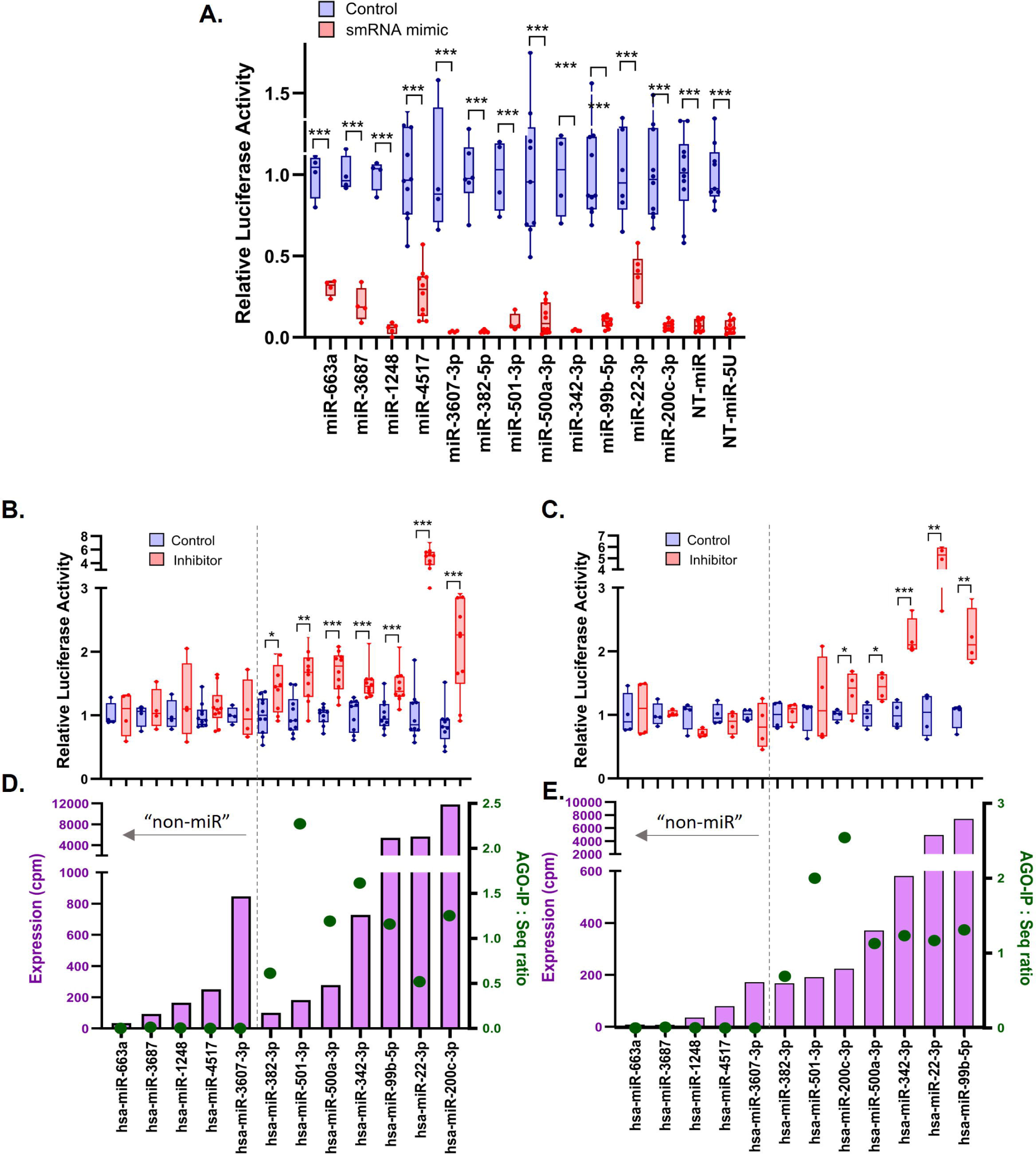
Non-AGO bound small RNAs at their endogenous levels do not repress luciferase reporters of fully complementary sequence. (A) Relative activity of fully complementary luciferase reporters co-transfected with their specific miRNA mimics (A, in mesHMLE cells) or inhibitors (B,C, in HMLE and mesHMLE cells respectively). Assays were performed with multiple replication with the values of independent biological replicates shown as dots. Relative luciferase activities were determined 48 h post-transfection. Pairwise t tests (comparing the mimic or inhibitor relative to the corresponding negative control) are shown: *p<0.05, **p<0.01, ***p<0.001. (D,E) Read counts (cpm) of RNAs from the whole cell sequencing (bars) in both D, HMLE and E, mesHMLE cells. Green dots indicate relative AGO binding efficiency. All miRNAs to the right of the dotted line are true, well established miRNAs and included as positive controls.

### Additional criteria for recognising non-miRs

Because AGO binding data may not be readily available for some small RNAs that have not been otherwise validated as miRNAs, we asked whether there are other features common to non-miRs that may aid their recognition, such as subcellular location, RNA length or 5’/3’ heterogeneity.

### Cytoplasmic versus nuclear location

Canonical miRNA processing concludes in the cytoplasm and it is within the cytoplasm that RISC resides and operates to post-transcriptionally silence genes. Other small RNAs can have different sub-cellular distributions; for example small nuclear and small nucleolar RNAs (snRNAs and snoRNAs) are nuclear. We noticed that for at least some of the non-AGO binding miRNAs, their genomic location overlaps with snoRNAs (Fig. 3A), suggesting the possibility that degraded products of larger RNAs may be mis-annotated as miRNAs. To investigate further, we isolated nuclear and cytoplasmic compartments from both HMLE and mesHMLE cells. Successful fractionation was confirmed by qPCR of known nuclear (NEAT1) and cytoplasmic (RPL30) RNAs and through the relative abundance of small RNA fragments derived from nuclear (snoRNA, snRNA) and cytoplasmic (tRNA) RNAs within our high-throughput sequencing libraries (Supplementary Figure S3). Due to different normalisation between cell compartments we cannot quantitatively compare miRNA abundance between nuclear and cytoplasmic fractions. However, it can still be seen that canonical miRNAs are readily detectable in both fractions, whereas the non-AGO binding “non-miRs” were almost exclusively nuclear in both HMLE and mesHMLE cells (Fig. 3B). Thus, nuclear localisation can aid recognition of non-miRs.

**Figure 3.**
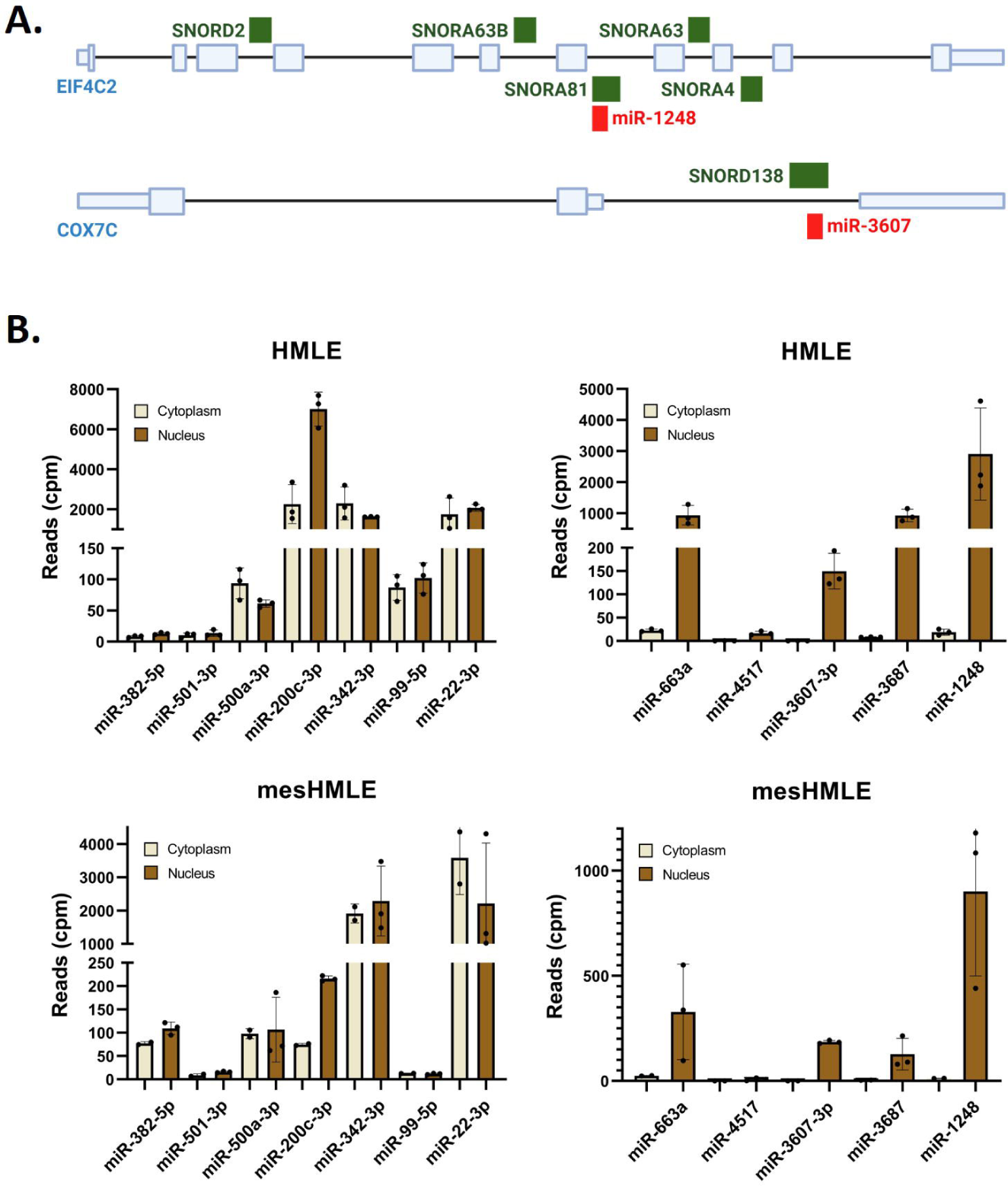
Annotated miRNAs that do not bind AGO and are enriched within nuclei. (A) Overlapping annotation of snoRNAs and miRNAs within the EIF4C2 and COX7C genes. (B) Read counts of individual miRNAs within the nuclei and cytoplasm of HMLE and mesHMLE cells, with the y-axis indicating read counts per million. The same small RNAs shown in Figure 1 are represented, including both true miRNAs (left panels) and non-miRs (right panels).

### Length heterogeneity

Authentic miRNAs typically have well defined 5’ ends and only a modest degree of length heterogeneity [10, 33–35]. To assess these features for non-miRs, we tabulated the start and end, and the length distribution, of RNA sequencing reads for several non-miRs, in comparison to two well-characterised authentic miRNAs, miR-22 and miR-200c (Fig. 4). MiR-22-3p and miR-200c-3p have features typical of functional miRNAs: i) AGO association, ii) cytoplasmic abundance, iii) length close to 22 nt and iv) high 5’ homogeneity. In contrast, each of the non-miRs tested, in addition to nuclear localisation and lack of AGO association, exhibit extended length (often >25nt) and in the cases of miR-663a, miR-1248 and miR-3687, highly variable 5’ ends (Figure 4). Looking broadly across miRNAs sequenced at sufficient depth in HMLE and/or mesHMLE samples, non-miRs (designated on the basis of no AGO binding) are far more likely to have a high proportion of reads >25nt in length and heterogeneity in 5’ ends, than are “true” miRNAs and “marginal” miRNAs, categorised on the basis of high and modest AGO binding respectively (Supplementary Figure 4, Supplementary Table 3). Taken together, examining the size and heterogeneity of small RNA reads can aid identification of non-miRs, even if additional information such as AGO interaction or subcellular localisation is not available.

**Figure 4.**
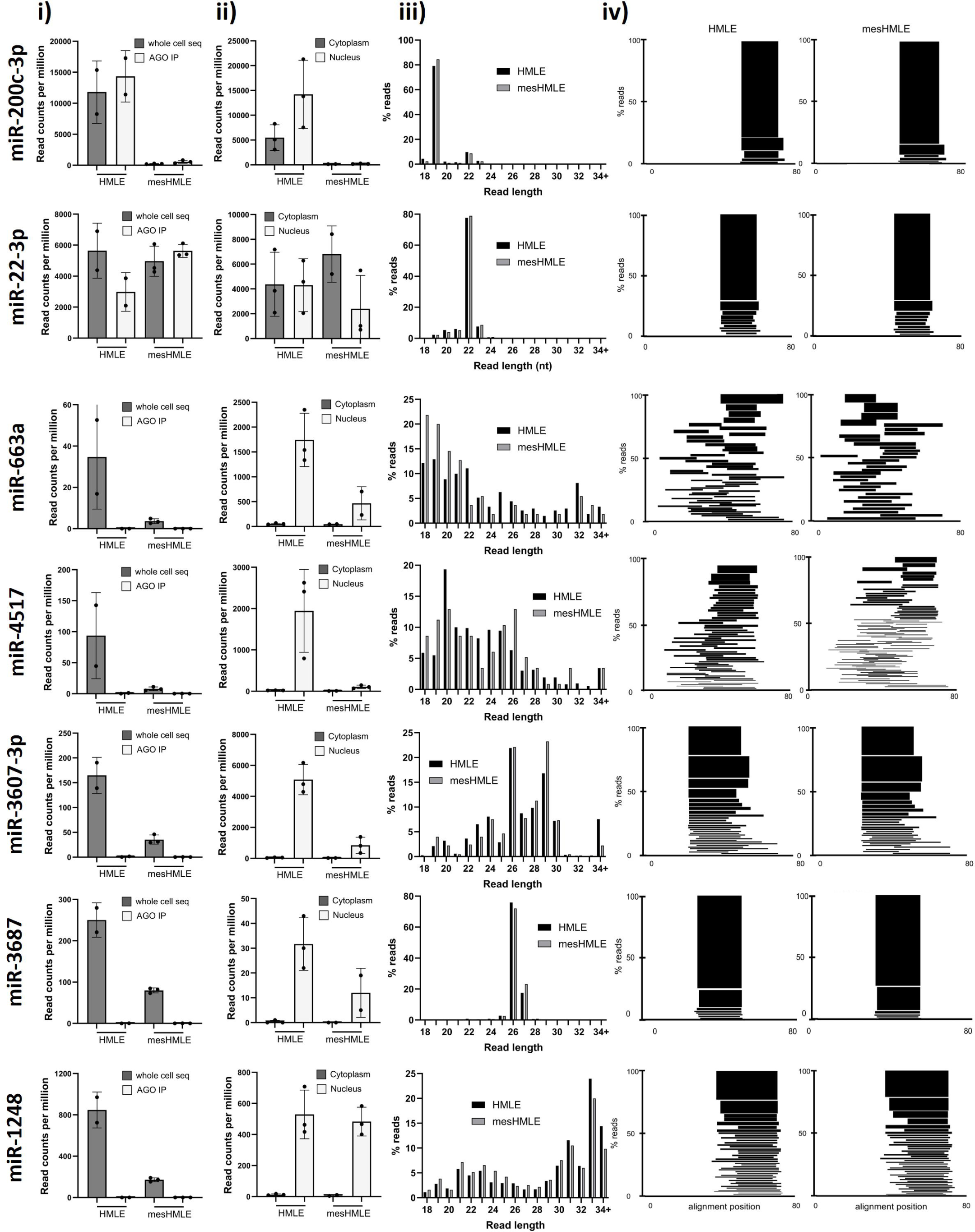
Small RNAs that do not associate with Argonaute display other characteristics atypical of genuine microRNAs. For a series of (A-B) well established miRNAs (miR-22-3p, -200c-3p) and (C-G) falsely annotated non-miRs, read counts in the (i) total RNA and AGO co-precipitate, (ii) nuclear and cytoplasmic fractions, (iii) size profile and (iv) distribution of mapped reads are shown. In iv), all sequencing reads mapping to that locus (representing >0.5% of all reads for that miRNA) are shown. The height of the bar represents the percentage of mapped reads. All alignments are across an 80nt window with the 5’ position relative to the start of the annotated pri-miR.

## Discussion

The work we present here represents the first investigation of which we are aware that examines AGO-association, a key feature of miRNA functionality, as a means to separate genuine and falsely annotated miRNAs. In so doing, we identify 27 “non-miRs” expressed in an EMT context, and note many hundreds of publications in which they feature or are a focus of. We expect however this is a considerable underestimate of the problem given we can only assess miRNAs that are expressed above a certain level, and the majority of miRbase-listed miRNAs do not meet this expression threshold (either here, or in most other contexts). We strongly suspect there will be other, likely hundreds, of non-miRs that have been given miRNA status, but that do not bind AGO, or do not achieve a level of expression sufficient for endogenous function. The need to include consideration of the relative cellular levels of a miRNA and proposed target, along with other issues that should be considered in assigning functions to miRNAs, has been well described by Kilikevicius et al. [36].

With this in mind, and given the issues with the interpretation of miRNA over-expression experiments we describe, we urge the cancer research community to utilise well controlled miRNA-inhibition experiments as a first-line of evidence when embarking on new investigations. Ideally, AGO co-precipitation experiments would also be conducted to establish AGO association but even if they never are, experiments involving miRNA inhibitors are an initial first step to establish if an endogenous miRNAs is both genuine and functionally significant. In our experience, the increased difficulty of getting miRNA-inhibition experiments to “work” – as opposed to over-expression experiments where function is relatively easily “demonstrated” – is not a methodological hurdle, but rather reflects an emerging understanding that many miRNAs are either falsely annotated RNA degradation products that do not bind AGO, and/or are simply expressed at insufficient levels for detectable function. Interestingly, most of the non-miRs are only expressed across cell lines and tissues at comparatively low levels (Supplementary Figure 5), meaning that even if they are capable of binding AGO or displaying other features consistent with miRNA functionality, expression level alone may preclude endogenous impact. This is yet another reason that establishing endogenous functionality as a first principal, achieved through the early implementation of miRNA inhibition, would vastly improve the cancer field which is plagued by overstated claims of miRNA function.

While there is recognition that the true number of human miRNAs may be much less than the several thousand that have been given the label “miRNA” [10], papers continue to be published that mistakenly ascribe miRNA functions to non-miRs. For example, in the past two years there have been 20 papers, including some in Q1 journals, reporting effects of miR-663a or miR-663b, which we show here are non-miRs. This may be in part due to a lack of clear guidelines for recognising which of the previously annotated miRNAs are actually not miRNAs, although the MirGeneDB database is a good starting point to check that a miRNA in question has the length and precursor features expected of a miRNA [37–41]. We show here those criteria correlate well with presence of the miRNA in the RISC complex in vivo, and ability of the endogenous miRNA to inhibit reporter expression.

While most miRNAs are produced by the canonical pathway involving precursor cleavage by Drosha and Dicer, there are exceptions, making the binding to AGO, rather than mode of biogenesis or structural features required for processing, a more useful criterion for miRNA status. For example, the production of miR-451 is independent of Dicer but requires AGO2 cleavage, followed by additional nucleotide trimming [42–44]. AGO2 is also required to cleave the passenger strand during the Drosha and Dicer-dependent production of miR-486-5p [45]. Drosha-independent, Dicer dependent miRNAs also exist including 5’-capped miRNAs, whose 5’ end corresponds to the transcription start site [46], and miRtrons whose non-canonical biogenesis requires the spliceosome [47, 48]. Additionally, at least some larger ncRNAs also encode genuine, AGO-binding smRNAs that act in the miRNA pathway, so that overlapping genomic annotations between a miRNA and another class of non-coding RNA are not sufficient evidence to discount miRNA status. These can be generated from snoRNAs [49–52], rRNAs [53] and tRNAs [54–58], in a typically Dicer-dependent manner [50, 55, 58].

Uncertainty may also arise from small RNAs that fulfil some, but not all, criteria one would expect of a miRNA. For example, if a small RNA interacts with AGO poorly, it may still be capable of suppressing target genes, but may do so with less efficiency than one would anticipate from the degree to which it is expressed. This may come about because it is unlikely a new miRNA would emerge from a single evolutionary event, leading to so called “transitional miRNAs”; small RNAs in the process of evolving miRNA-like activity, but whose gene silencing roles remain limited [34, 35]. In the cases of miR-221 and miR-148 for example, the degree to which they co-precipitate with AGO is only modest, though both are highly expressed and thus represent significant fractions of the AGO-abound miRNAome. The significance of other, more lowly expressed RNAs that co-precipitate with similarly modest efficiency, are more questionable. In Supplementary Table 3, we therefore include “marginal” miRNAs (in addition to genuine and falsely annotated listings), that possess some, but not all, features one would expect of efficient, evolutionarily optimised miRNAs.

Because it is known that not all miRBase entries are genuine, in its latest release (v22) a text-mining approach has been employed to improve confidence in the functionality of listed miRNAs, scanning publications for miRNA name-containing sentences and attributing positive scores for the co-existence of terms related to function (such as “expression”, “target”, “regulate” or “inhibit”) [15]. This however is problematic if there are small RNAs that do not endogenously function as miRNAs, but are nevertheless reported as miRNAs based upon exogenous expression and/or an incorrect interpretation of experiments involving single or poorly controlled miRNA inhibitors. Importantly, in Figure 2 we demonstrate that even miRNA-sized sequences not encoded within the genome can still behave as though they are miRNAs when transiently expressed. The existence of reports of miRNAs in which apparent function is shown through over-expression then reaffirms their listing in miRBase, which in turn makes it more likely that future researchers will perform similar experiments. This circular logic, we believe, is responsible for thousands of publications involving spurious “miRNAs”, including in the cancer field where we believe this false reporting underlies thousands of cancer publications.

**Table 1.** Chemicals, biological reagents and computational resources.

| Reagents/kits | Resource | Catalogue number/ reference |
| --- | --- | --- |
| 10x PBS | Gibco | 70011044 |
| 2x Universal PCR Master Mix | Applied Biosystems | 4324018 |
| Bioanalyzer small RNA assay | Agilent | 5067-1548 |
| NEXTflex Small RNA-Seq Kit v3 | Perkin Elmer | NOVA-5132-05 |
| DMEM:F12 media | Gibco | 11320033 |
| Dual-Luciferase® Reporter | Promega | E1980 |
| Ethylenediaminetetraacetic acid | Univar Solutions | A180 |
| Ethanol | Univar Solutions | AJA214 |
| Glycerol | Univar Solutions | 242 |
| HuMEC Basal Serum-Free Medium | Gibco | 12753018 |
| HuMEC Supplement Kit | Gibco | 12755013 |
| Hydrocortisone | Sigma Aldrich | H0888 |
| Insulin | Actrapid | N/A |
| Lipofectamine™ 2000 | Invitrogen | 11668019 |
| Magnesium chloride | Merck | 7791186 |
| Multiplex TaqMan miRNA Reverse<br>Transcription Kit | Applied Biosystems | 4366596 |
| Nanodrop Qubit HS RNA | Invitrogen | Q32852 |
| NotI-HF Restriction enzyme | NewEngland BioLabs | R3189S |
| NP-40 | IGEPAL, # CA-630 | CA-630 |
| NP-40 alternative | Calbiochem | 492016 |
| Pan-anti-Ago antibody | 4F9 | [19] |
| Phenol | Ambion | AM9712 |
| Pierce protein-L magnetic beads | Thermo Scientific | 88849 |
| Protease inhibitors | Roche | 4693159001 |
| Proteinase K | Invitrogen | 25530049 |
| QIAseq miRNA kit | Qiagen | 331502 |
| QuantiTect Reverse Transcription | Qiagen | 205311 |
| QuantiTect SYBR Green PCR Kit | Qiagen | 204141 |
| Recombinant Human EGF Protein | R&D systems | RDS236EG01M |
| Recombinant Human TGF-beta 1 | R&D systems | RDS240B002 |
| RNAseOUT | Invitrogen | 10777019 |
| Sodium acetate | AnalaR | 10236 |
| Sodium chloride | Chem Supply | SA046 |
| Sodium deoxycholate | Sigma Aldrich | D6750 |
| Sodium dodecyl sulfate | Sigma Aldrich | 75746 |
| SUPERase-In | ThermoFisher | AM2694 |
| T4 DNA Ligase | NewEngland BioLabs | M0202S |
| Tris | Invitrogen | 15504020 |
| Trizol Reagent | Ambion | 15596018 |
| TurboDNase | Ambion | AM2238 |
| XhoI Restriction enzyme | NewEngland BioLabs | R0146S |
| **Cell lines/ plasmids** | **Resource** | **Catalogue number/reference** |
| HMLE cells | Weinberg lab | [17] |
| Fetal Bovine serum | Hyclone | SH300084 |
| PsiCHECK2 | Promega | C8021 |
| **Tools/ databases** | **Resource** |  |
| FastQC v0.11.9 | [21] |  |
| Cutadapt v2.8 | [22] |  |
| UMI-tools v1.0.1 | [23] |  |
| BWA v0.7.17 alignment algorithm | [24] |  |
| Integrative Genomics Viewer v2.8.0 | [25] |  |
| CalcNormFactors - edgeR package | [30] |  |
| GraphPad Prism | GraphPad Co. |  |
| GRCh37 annotation ensemble | [26] |  |
| miRbase | [27] |  |
| Human reference genome (hg19) | www.ncbi.nlm.nih.gov/assembly/GCF_000001405.13/ |  |

## Acknowledgements

This work was supported by funding from the Australian Research Council (FT190100544, DP190103333), the Cancer Council Beat Cancer Project (PRF2518) and the Worldwide Cancer Research Foundation (WCR-19-0300).

K.A.P. was supported by the Royal Adelaide Hospital Research Committee Florey Fellowship.

## Author contributions

AO performed cell fractionation, cloned reporter constructs and performed most of the luciferase assays. Additional luciferases were conducted by MM. BKD and AGB performed the HMLE and mesHMLE AGO-co-precipitations. NIW led the bioinformatic work, supported by JMB and KAP. PAG and GJG provided oversight, intellectual contribution and critical editing. CPB directed the project, led data analysis, and along with GJG wrote the manuscript.

## Declaration of interests

The authors have no competing interests to declare. Data Availability

Raw fastq files for the small RNA-seq and AGO-IP data are available from the Gene Expression Omnibus under the accessions GSE184533 and GSE184534, respectively. Raw fastq files and processed gene expression data for nuclear and cytoplasmic fractionation data are available under the accession GSE183863.

## References

1 Kobayashi H, Tomari Y. RISC assembly: Coordination between small RNAs and Argonaute proteins. Biochimica et biophysica acta 2016; 1859: 71–81.

2 Nakanishi K. Anatomy of four human Argonaute proteins. Nucleic Acids Res 2022; 50: 6618–6638.

3 Bartel DP. Metazoan MicroRNAs. Cell 2018; 173: 20–51.

4 Bracken CP, Scott HS, Goodall GJ. A network-biology perspective of microRNA function and dysfunction in cancer. Nat Rev Genet 2016; 17: 719–732.

5 Nemeth K, Bayraktar R, Ferracin M, Calin GA. Non-coding RNAs in disease: from mechanisms to therapeutics. Nat Rev Genet 2024; 25: 211–232.

6 Bracken CP, Goodall GJ, Gregory PA. RNA regulatory mechanisms controlling TGF-beta signaling and EMT in cancer. Semin Cancer Biol 2024; 102–103: 4-16.

7 Cursons J, Pillman KA, Scheer KG, Gregory PA, Foroutan M, Hediyeh-Zadeh S et al. Combinatorial Targeting by MicroRNAs Co-ordinates Post-transcriptional Control of EMT. Cell Syst 2018; 7: 77–91 e77.

8 Yang J, Antin P, Berx G, Blanpain C, Brabletz T, Bronner M et al. Guidelines and definitions for research on epithelial-mesenchymal transition. Nat Rev Mol Cell Biol 2020; 21: 341–352.

9 Alles J, Fehlmann T, Fischer U, Backes C, Galata V, Minet M et al. An estimate of the total number of true human miRNAs. Nucleic Acids Res 2019; 47: 3353–3364.

10 Fromm B, Zhong X, Tarbier M, Friedlander MR, Hackenberg M. The limits of human microRNA annotation have been met. RNA 2022; 28: 781–785.

11 Hansen TB, Kjems J, Bramsen JB. Enhancing miRNA annotation confidence in miRBase by continuous cross dataset analysis. RNA Biol 2011; 8: 378–383.

12 Langenberger DB, S.; Hertel, J.; Hoffmann, S.; Tafer, H.; Stadler, P.F. MicroRNA or Not MicroRNA? Advances in Bioinformatics and Computational Biology 2011: 1–9.

13 Schopman NC, Heynen S, Haasnoot J, Berkhout B. A miRNA-tRNA mix-up: tRNA origin of proposed miRNA. RNA Biol 2010; 7: 573–576.

14 Thomson DW, Pillman KA, Anderson ML, Lawrence DM, Toubia J, Goodall GJ et al. Assessing the gene regulatory properties of Argonaute-bound small RNAs of diverse genomic origin. Nucleic Acids Res 2015; 43: 470–481.

15 Kozomara A, Birgaoanu M, Griffiths-Jones S. miRBase: from microRNA sequences to function. Nucleic acids research 2019; 47: D155–d162.

16 Fromm B, Domanska D, Hoye E, Ovchinnikov V, Kang W, Aparicio-Puerta E et al. MirGeneDB 2.0: the metazoan microRNA complement. Nucleic Acids Res 2020; 48: D1172.

17 Mani SA, Guo W, Liao MJ, Eaton EN, Ayyanan A, Zhou AY et al. The epithelial-mesenchymal transition generates cells with properties of stem cells. Cell 2008; 133: 704–715.

18 Gagnon KT, Li L, Janowski BA, Corey DR. Analysis of nuclear RNA interference in human cells by subcellular fractionation and Argonaute loading. Nature protocols 2014; 9: 2045–2060.

19 Ikeda K, Satoh M, Pauley KM, Fritzler MJ, Reeves WH, Chan EK. Detection of the argonaute protein Ago2 and microRNAs in the RNA induced silencing complex (RISC) using a monoclonal antibody. Journal of immunological methods 2006; 317: 38–44.

20 Jensen KB, Darnell RB. CLIP: crosslinking and immunoprecipitation of in vivo RNA targets of RNA-binding proteins. Methods in molecular biology (Clifton, NJ) 2008; 488: 85–98.

21 Andrews S. FastQC: A quality control tool for high throughput sequence data. https://wwwbioinformaticsbabrahamacuk/projects/fastqc/ 2010.

22 Martin M. Cutadapt removes adapter sequences from high-throughput sequencing reads. EMBnetjournal 2011; 17(1): 10–12.

23 Smith T, Heger A, Sudbery I. UMI-tools: modeling sequencing errors in Unique Molecular Identifiers to improve quantification accuracy. Genome Res 2017; 27: 491–499.

24 Li H, Durbin R. Fast and accurate long-read alignment with Burrows-Wheeler transform. Bioinformatics 2010; 26: 589–595.

25 Robinson JT, Thorvaldsdóttir H, Winckler W, Guttman M, Lander ES, Getz G et al. Integrative genomics viewer. Nature biotechnology 2011; 29: 24–26.

26 Howe KL, Achuthan P, Allen J, Allen J, Alvarez-Jarreta J, Amode MR, et al. Ensembl 2021. Nucleic Acids Res 2021; 49: D884–D891.

27 Griffiths-Jones S. The microRNA Registry. Nucleic Acids Res 2004; 32: D109–111.

28 Saunders K, Bert AG, Dredge BK, Toubia J, Gregory PA, Pillman KA et al. Insufficiently complex unique-molecular identifiers (UMIs) distort small RNA sequencing. Sci Rep 2020; 10: 14593.

29 Robinson MD, Oshlack A. A scaling normalization method for differential expression analysis of RNA-seq data. Genome Biol 2010; 11: R25.

30 Robinson MD, McCarthy DJ, Smyth GK. edgeR: a Bioconductor package for differential expression analysis of digital gene expression data. Bioinformatics 2010; 26: 139–140.

31 Migault M, Sapkota S, Bracken CP. Transcriptional and post-transcriptional control of epithelial-mesenchymal plasticity: why so many regulators? Cell Mol Life Sci 2022; 79: 182.

32 Petri R, Brattas PL, Sharma Y, Jonsson ME, Pircs K, Bengzon J et al. LINE-2 transposable elements are a source of functional human microRNAs and target sites. PLoS Genet 2019; 15: e1008036.

33 Chiang HR, Schoenfeld LW, Ruby JG, Auyeung VC, Spies N, Baek D et al. Mammalian microRNAs: experimental evaluation of novel and previously annotated genes. Genes Dev 2010; 24: 992–1009.

34 Ruby JG, Jan C, Player C, Axtell MJ, Lee W, Nusbaum C et al. Large-scale sequencing reveals 21U-RNAs and additional microRNAs and endogenous siRNAs in C. elegans. Cell 2006; 127: 1193–1207.

35 Yu F, Pillman KA, Neilsen CT, Toubia J, Lawrence DM, Tsykin A et al. Naturally existing isoforms of miR-222 have distinct functions. Nucleic Acids Res 2017; 45: 11371–11385.

36 Kilikevicius A, Meister G, Corey DR. Reexamining assumptions about miRNA-guided gene silencing. Nucleic Acids Res 2022; 50: 617–634.

37 Baek SC, Kim B, Jang H, Kim K, Park IS, Min DH et al. Structural atlas of human primary microRNAs generated by SHAPE-MaP. Mol Cell 2024; 84: 1158–1172 e1156.

38 Fromm B, Hoye E, Domanska D, Zhong X, Aparicio-Puerta E, Ovchinnikov V et al. MirGeneDB 2.1: toward a complete sampling of all major animal phyla. Nucleic Acids Res 2022; 50: D204–D210.

39 Kim K, Baek SC, Lee YY, Bastiaanssen C, Kim J, Kim H et al. A quantitative map of human primary microRNA processing sites. Mol Cell 2021; 81: 3422–3439 e3411.

40 Krol J, Sobczak K, Wilczynska U, Drath M, Jasinska A, Kaczynska D et al. Structural features of microRNA (miRNA) precursors and their relevance to miRNA biogenesis and small interfering RNA/short hairpin RNA design. J Biol Chem 2004; 279: 42230–42239.

41 Starega-Roslan J, Krol J, Koscianska E, Kozlowski P, Szlachcic WJ, Sobczak K et al. Structural basis of microRNA length variety. Nucleic Acids Res 2011; 39: 257–268.

42 Cheloufi S, Dos Santos CO, Chong MM, Hannon GJ. A dicer-independent miRNA biogenesis pathway that requires Ago catalysis. Nature 2010; 465: 584–589.

43 Cifuentes D, Xue H, Taylor DW, Patnode H, Mishima Y, Cheloufi S et al. A novel miRNA processing pathway independent of Dicer requires Argonaute2 catalytic activity. Science 2010; 328: 1694–1698.

44 Yang JS, Maurin T, Robine N, Rasmussen KD, Jeffrey KL, Chandwani R et al. Conserved vertebrate mir-451 provides a platform for Dicer-independent, Ago2-mediated microRNA biogenesis. Proc Natl Acad Sci U S A 2010; 107: 15163–15168.

45 Jee D, Yang JS, Park SM, Farmer DT, Wen J, Chou T et al. Dual Strategies for Argonaute2-Mediated Biogenesis of Erythroid miRNAs Underlie Conserved Requirements for Slicing in Mammals. Mol Cell 2018; 69: 265–278 e266.

46 Xie M, Li M, Vilborg A, Lee N, Shu MD, Yartseva V et al. Mammalian 5’-capped microRNA precursors that generate a single microRNA. Cell 2013; 155: 1568–1580.

47 Okamura K, Hagen JW, Duan H, Tyler DM, Lai EC. The mirtron pathway generates microRNA-class regulatory RNAs in Drosophila. Cell 2007; 130: 89–100.

48 Ruby JG, Jan CH, Bartel DP. Intronic microRNA precursors that bypass Drosha processing. Nature 2007; 448: 83–86.

49 Brameier M, Herwig A, Reinhardt R, Walter L, Gruber J. Human box C/D snoRNAs with miRNA like functions: expanding the range of regulatory RNAs. Nucleic Acids Res 2011; 39: 675–686.

50 Ender C, Krek A, Friedlander MR, Beitzinger M, Weinmann L, Chen W et al. A human snoRNA with microRNA-like functions. Mol Cell 2008; 32: 519–528.

51 Patterson DG, Roberts JT, King VM, Houserova D, Barnhill EC, Crucello A et al. Human snoRNA-93 is processed into a microRNA-like RNA that promotes breast cancer cell invasion. NPJ Breast Cancer 2017; 3: 25.

52 Yu F, Bracken CP, Pillman KA, Lawrence DM, Goodall GJ, Callen DF et al. p53 Represses the Oncogenic Sno-MiR-28 Derived from a SnoRNA. PLoS One 2015; 10: e0129190.

53 Guan L, Grigoriev A. Computational meta-analysis of ribosomal RNA fragments: potential targets and interaction mechanisms. Nucleic Acids Res 2021; 49: 4085–4103.

54 Guan L, Karaiskos S, Grigoriev A. Inferring targeting modes of Argonaute-loaded tRNA fragments. RNA Biol 2020; 17: 1070–1080.

55 Hasler D, Lehmann G, Murakawa Y, Klironomos F, Jakob L, Grasser FA et al. The Lupus Autoantigen La Prevents Mis-channeling of tRNA Fragments into the Human MicroRNA Pathway. Mol Cell 2016; 63: 110–124.

56 Kumar P, Anaya J, Mudunuri SB, Dutta A. Meta-analysis of tRNA derived RNA fragments reveals that they are evolutionarily conserved and associate with AGO proteins to recognize specific RNA targets. BMC Biol 2014; 12: 78.

57 Kuscu C, Kumar P, Kiran M, Su Z, Malik A, Dutta A. tRNA fragments (tRFs) guide Ago to regulate gene expression post-transcriptionally in a Dicer-independent manner. RNA 2018; 24: 1093–1105.

58 Maute RL, Schneider C, Sumazin P, Holmes A, Califano A, Basso K et al. tRNA-derived microRNA modulates proliferation and the DNA damage response and is down-regulated in B cell lymphoma. Proc Natl Acad Sci U S A 2013; 110: 1404–1409.

