## Supplementary Data for "Chasing non-existent “microRNAs” in cancer"

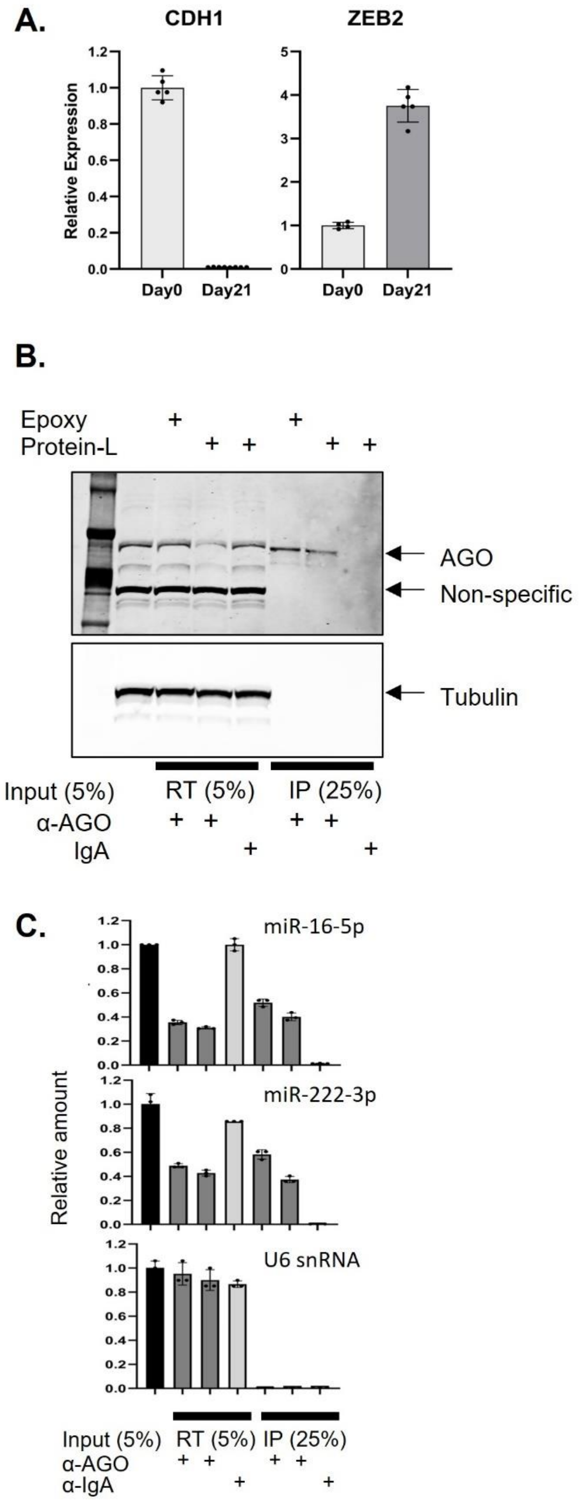


**Supplementary Figure 1. MicroRNAs are specifically co-precipitated with Argonaute**

(A) Successful establishment of a mesenchymal cell line ("mesHMLEs") derived from HMLE cells after 21 days of TGF-β treatment is indicated by the relative expression levels of epithelial (CDH1) and mesenchymal (ZEB2) markers. (B) AGO proteins are effectively and specifically immunoprecipitated by a pan-AGO antibody using either Epoxy or Protein-L magnetic beads. R/T = run-through; IP = Immunoprecipitate (C) TaqMan qPCR of miRNA & control U6 show specificity of miRNAs co-precipitated with AGO.

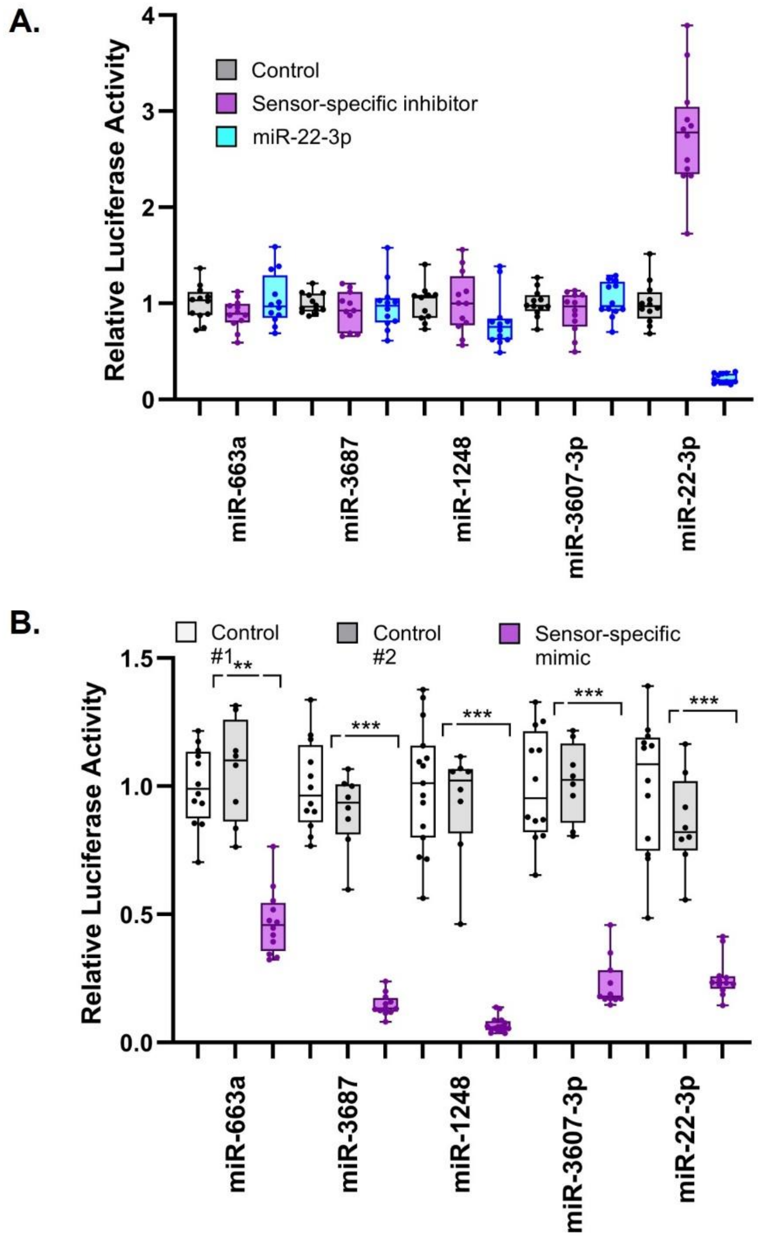


**Supplementary Figure 2. Additional controls for luciferase reporter assays**

Relative luciferase activity of fully complementary luciferase reporters co-transfected with either (A) their specific miRNA mimics or two independent miRNA controls or (B) miRNA inhibitors that are specific to the either the reporter, or to miR-22-3p. A miR-22-3p mimic is also co-transfected to further indicate specificity of target suppression. Independent replicates are indicated. Pairwise t tests (comparing the mimic or inhibitor relative to the corresponding negative control) are shown: *p<0.05, **p<0.01, ***p<0.001.

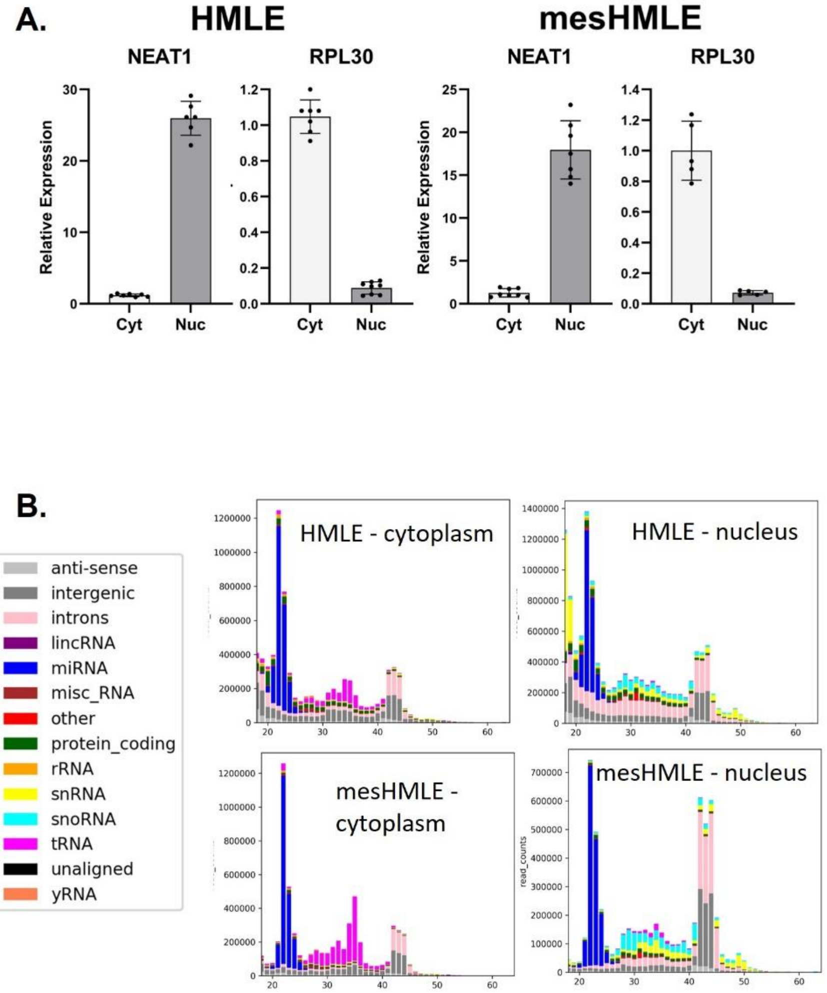


**Supplementary Figure 3. Subcellular fractionation**

(A) HMLE and mesHMLE cells were separated into cytoplasmic and nuclear fractions and subjected to small RNA sequencing. The purity of subcellular fractionation was confirmed by measuring NEAT1 expression (nuclear marker) and RPL30 expression (cytoplasmic marker) by qPCR. Results are shown as mean ± SD of three biological replicates. B) Successful fractionation of the cells was further indicated by exclusive nuclear presence of snoRNAs (light blue bars) and snRNAs (yellow bars) and cytoplasmic presence of tRNAs (dark pink bars). The y and x-axes indicate total sequencing read counts and length of the associated small RNA, respectively.

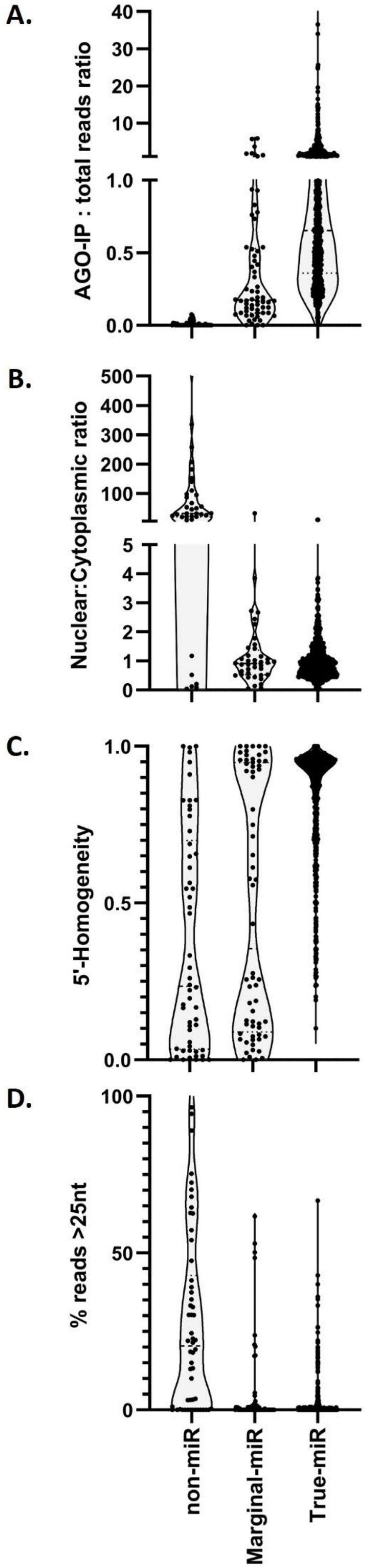


**Supplementary Figure 4. Nuclear localisation, 5'-heterogeneity and read length correlate with, but are not definitive markers of, falsely-annotated miRNAs**

A) Relative AGO binding efficiency, B) nuclear : cytoplasmic distribution, C) 5'-homogeneity, being the fraction of sequencing reads that have the most common 5' end and D) proportion of sequencing reads above 25 nt for both HMLE and mesHMLE cells. MiRNAs are divided into: non-miRs (the 27 miRNAs listed in Figure 1), true miRs and marginal miRs (listed in Supplementary Table 3). In each case, for representation all sequences must be detected in HMLE and/or mesHMLE cells at >5 cpm.

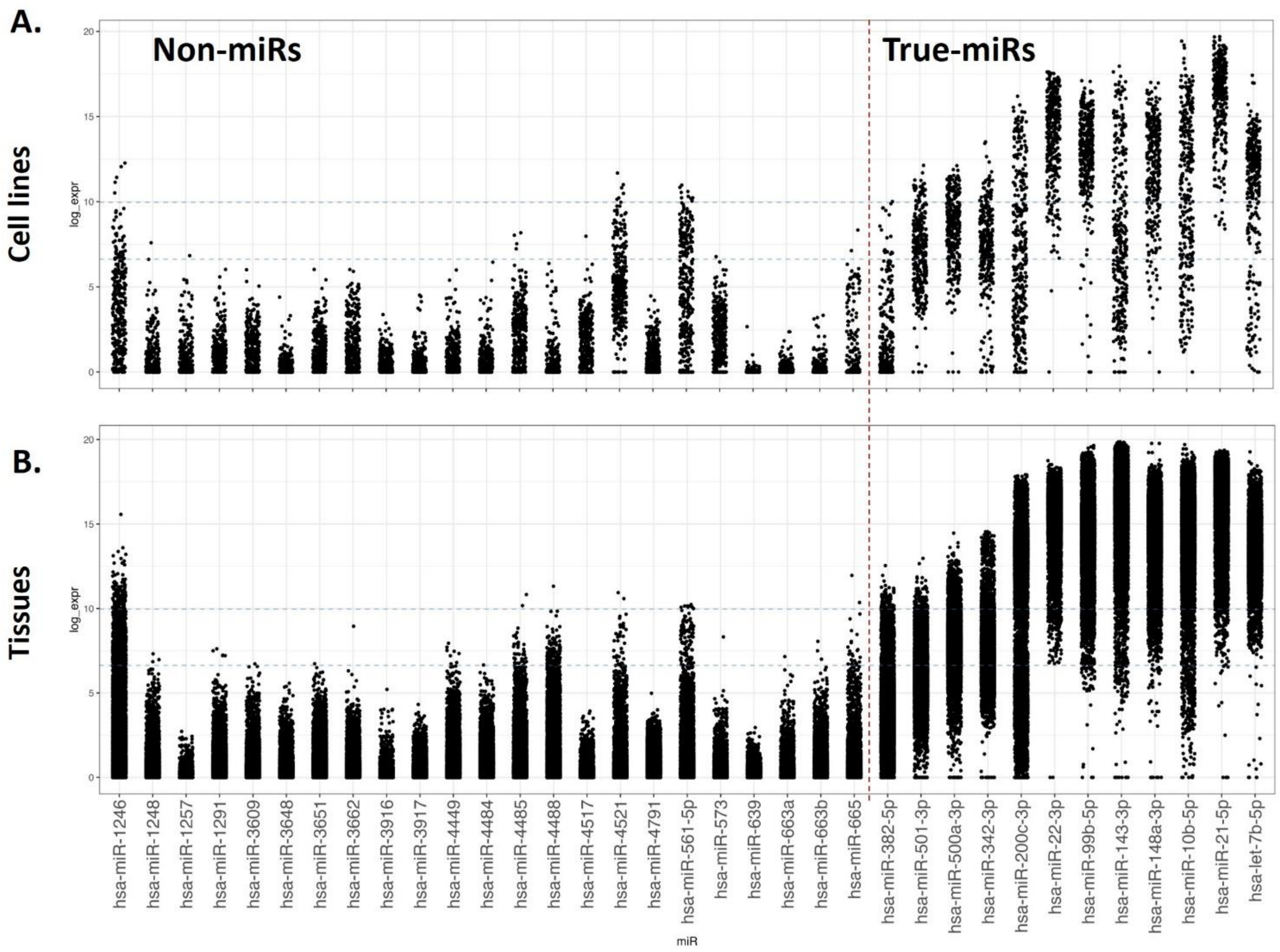


**Supplementary Figure 5. Most non-miRs are expressed across cell lines and tissues at very low levels**

Maximal miRNA expression for each non-miR (left) and a series of true miRNAs (right) across A) 333 cell lines and B) 10,716 tissue samples from >50 diverse cancer types (data obtained from [59]). Each dot represents the maximal expression of that miRNA in the total RNA sequencing (cpm). A log10 scale is employed to aid visualisation.

| Corresponding miRNA | Primer sequence (5′-3′) |
| --- | --- |
| miR-663a | 5′- TCGAGATTTAAATGCGGTCCCGCGGCGCCCCGCCTGC -3′ |
| | 5′- GGCCGCAGGCGGGGCGCCGCGGGACCGCATTTAAATC -3′ |
| miR-4517 | 5′- TCGAGATTTAAATCTCAGCTGTGAGTTTCATCATATTTGC -3′ |
| | 5′- GGCCGCAAATATGATGAAACTCACAGCTGAGATTTAAATC -3′ |
| miR-3607-3p | 5′- TCGAGATTTAAATCATCAGAAAGCGTTTACAGTGC -3′ |
| | 5′- GGCCGCACTGTAAACGCTTTCTGATGATTTAAATC -3′ |
| miR-3687 | 5′- TCGAGATTTAAATACGTCGCACGAACGCCTGTCCGGGGC- -3′ |
| | 5′- GGCCGCCCCGGACAGGCGTTCGTGCGACGTATTTAAATC -3′ |
| miR-1248 | 5′- TCGAGATTTAAATTTTAGCACAGTGCTTATACAAGAAGGTGC -3′ |
| | 5′- GGCCGCACCTTCTTGTATAAGCACTGTGCTAAAATTTAAATC -3′ |
| miR-382-5p | 5′- TCGAGATTTAAATCGAATCCACCACGAACAACTTCGC -3′ |
| | 5′- GGCCGCGAAGTTGTTCGTGGTGGATTCGATTTAAATC -3′ |
| miR-500a-3p | 5′- TCGAGATTTAAATCAGAATCCTTGCCCAGGTGCATGC -3′ |
| | 5′- GGCCGCATGCACCTGGGCAAGGATTCTGATTTAAATC -3′ |
| miR-501-3p | 5′- TCGAGATTTAAATAGAATCCTTGCCCGGGTGCATTGC -3′ |
| | 5′- GGCCGCAATGCACCCGGGCAAGGATTCTATTTAAATC -3′ |
| miR-99a-3p | 5′- TCGAGATTTAAATCAGACCCATGAAGCGAGCTTGGC -3′ |
| | 5′- GGCCGCCAAGCTCGCTTCTATGGGTCTGATTTAAATC -3′ |
| miR-342-3p | 5′- TCGAGATTTAAATACGGGTGCGATTTCTGTGTGAGAGC -3′ |
| | 5′- GGCCGCTCTCACACAGAAATCGCACCCGTATTTAAATC -3′ |
| miR-22-3p | 5′- TCGAGATTTAAATACAGTTCTTCAACTGGCAGCTTGC -3′ |
| | 5′- GGCCGCAAGCTGCCAGTTGAAGAACTGTATTTAAATC -3′ |
| miR-200c-3p | 5′- TCGAGATTTAAATTCCATCATTACCCGGCAGTATTAGC -3′ |
| | 5′- GGCCGCTAATACTGCCGGGTAATGATGGAATTTAAATC -3′ |
| NT-miR | 5′- TCGAGATTTAAATTGGCGTAACTCCGCGTATGGGC -3′ |
| | 5′- GGCCGCCCATACGCGGAGTTACGCCAATTTAAATC -3′ |
| NT-miR-5U | 5′- TCGAGATTTAAATGTACGTGACACGTTCGGAGAAGC -3′ |
| | 5′- GGCCGCTTCTCCGAACGTGTCACGTACATTTAAATC -3′ |

**Supplementary Table 1. Primers annealed for psiCheck2 dual luciferase cloning**

| | | |
|---|---|---|
| miR-663a | Mimic sense: | 5′- AGGCGGGGCGCCGCGGGACCGC -3′ |
| | Mimic antisense: | 3′- UUUCCGCCCCGCGGCGCCCUGG -5′ |
| | Inhibitor: | 5′- GCGGUCCCGCGGCGCCCCGCCU -3′ |
| miR-4517 | Mimic sense: | 5′- AAAUAUGAUGAAACUCACAGCUGAG -3′ |
| | Mimic antisense: | 3′- UUUUUAUACUACUUUGAGUGUCGAC -5′ |
| | Inhibitor: | 5′- CTCAGCUGUGAGUUUCAUCAUAUUU -3′ |
| miR-3607-3p | Mimic sense: | 5′- ACUGUAAACGCUUUCUGAUG -3′ |
| | Mimic antisense: | 3′- UUUGACAUUUGCGAAAGACU -5′ |
| | Inhibitor: | 5′- CAUCAGAAAGCGUUUACAGU -3′ |
| miR-3687 | Mimic sense: | 5′- CCCGGACAGGCGUUCGUGCGACGU -3′ |
| | Mimic antisense: | 3′-UUGGGCCUGUCCGCAAGCACGCUG -5′ |
| | Inhibitor: | 5′- ACGUCGCACGAACGCCUGUCCGGG -3′ |
| miR-1248 | Mimic sense: | 5′- CCUUCUUGUAUAAGCACUGUGCUAAA -3′ |
| | Mimic antisense: | 3′- UUGGAAGAACAUAUUCGUGACACGAU -5′ |
| | Inhibitor: | 5′- UUUAGCACAGUGCUUAUACAAGAAGG -3′ |
| miR-382-5p | Mimic sense: | 5′- GAAGUUGUUCGUGGUGGAUUCG -3′ |
| | Mimic antisense: | 3′- UUCUUCAACAAGCACCACCUAA -5′ |
| | Inhibitor: | 5′- CGAAUCCACCACGAACAACUUC -3′ |
| miR-500a-3p | Mimic sense: | 5′- AUGCACCUGGGCAAGGAUUCUG -3′ |
| | Mimic antisense: | 3′- UUUACGUGGACCCGUUCCUAAG -5′ |
| | Inhibitor: | 5′- CAGAAUCCUUGCCCAGGUGCAU -3′ |
| miR-501-3p | Mimic sense: | 5′- AAUGCACCCGGGCAAGGAUUCU -3′ |
| | Mimic antisense: | 3′- UUUUACGUGGGCCCGUUCCUAA -5′ |
| | Inhibitor: | 5′- AGAAUCCUUGCCCGGGUGCAUU -3′ |
| miR-99a-3p | Mimic sense: | 5′- CAAGCUCGCUUCUAUGGGUCUG -3′ |
| | Mimic antisense: | 3′- UUGUUCGAGCGAAGAUACCCAG -5′ |
| | Inhibitor: | 5′- CAGACCCAUAGAAGCGAGCUUG -3′ |
| miR-342-3p | Mimic sense: | 5′- UCUCACACAGAAAUCGCACCCGU -3′ |
| | Mimic antisense: | 3′- UUAGAGUGUGUCUUUAGCGUGGG -5′ |
| | Inhibitor: | 5′- ACGGGUGCGAUUUCUGUGUAGA -3′ |
| miR-22-3p | Mimic sense: | 5′- AAGCUGCCAGUUGAAGAACUGU -3′ |
| | Mimic antisense: | 3′- UUUUCGACGGUCAACUUCUUGA -5′ |
| | Inhibitor: | 5′- ACAGUUCUUCAACUGGCAGCUU -3′ |
| miR-200c-3p | Mimic sense: | 5′- UAAUACUGCCGGGUAAUGAUGGA -3′ |
| | Mimic antisense: | 3′- UUAUUAUGACGGCCCAUUACUAC -5′ |
| | Inhibitor: | 5′- UCCAUCAUUACCCGGCAGUAUUA -3′ |
| NT-miR | Mimic sense: | 5′- CCAUACGCGGAGUUACGCCAC -3′ |
| | Mimic antisense: | 3′- UUGGUAUGCGCCUCAAUGCGG -5′ |
| NT-miR-5U | Mimic sense: | 5′- UUCUCCGAACGUGUCACGUAC -3′ |
| | Mimic antisense: | 3′- UUAAGAGGCUUGCACAGUGCA -5′ |

**Supplementary Table 2. microRNA mimics and inhibitor sequences (5’-3’)**

**Supplementary Table 3. Non-miRs, true miRs and marginal miRs**

(excel file)
